# Confinement size determines the architecture of Ran-induced microtubule networks

**DOI:** 10.1101/2020.09.01.277186

**Authors:** Ya Gai, Sagar Setru, Brian Cook, Howard A. Stone, Sabine Petry

## Abstract

The organization of microtubules (MTs) within cells is critical for its internal organization during interphase and mitosis. During mitotic spindle assembly, MTs are made and organized around chromosomes in a process regulated by RanGTP. The role of RanGTP has been explored in *Xenopus* egg extracts, which are not limited by a cell membrane. Here, we investigated whether cell-sized confinements affect the assembly of RanGTP-induced MT networks in *Xenopus* egg extracts. We used microfluidics to encapsulate extract within monodisperse extract-in-oil droplets. Importantly, we find that the architecture of Ran-induced MT networks depends on the droplet diameter and the Ran concentration, and differs from structures formed in bulk extract. Our results highlight that both MT nucleation and physical confinement play critical roles in determining the spatial organization of the MT cytoskeleton.

**STATEMENT OF SIGNIFICANCE:** During cell division, chromosomes are segregated by the mitotic spindle, whose framework consists of up to hundreds of thousands of microtubules (MTs). Spindle MTs are generated via several pathways, one of which is regulated by RanGTP. Yet, how Ran-induced MTs self-organize within cell-sized confinement remains unclear. This work reports unexpected architectures of Ran-induced MT networks confined in cell-sized droplets, which depend on the droplet diameter and the RanGTP concentration. Thus, MT nucleation and confinement together give rise to specific MT network architectures, which are otherwise not observed in unconfined assays. The findings provide a simple strategy to engineer the architectures of MT networks and could have direct implications in nucleation-controlled soft material processing.

## INTRODUCTION

In the course of a cell cycle, a variety of morphological events occur, which are guided by cytoskeletal filaments confined within the cell. A key event in the proliferation of eukaryotic cells is the assembly of the mitotic spindle, which ensures the equal partitioning of chromosomes during cell division (1). Central to spindle assembly is the formation and spatial organization of MTs, which occurs to a great extent from chromosomes via the protein RanGTP, which is the upstream factor of a pathway that is even more critical for spindles in the absence of centrosomes (2–7).

During spindle assembly, chromosomes are surrounded by a concentration gradient of RanGTP (1, 8), which is generated by the chromatin-bound guanine nucleotide exchange factor RCC1 and the GTPase-activating protein RanGAP residing in the cytoplasm (3, 5). RanGTP releases spindle-assembly factors (SAF) from importins, which are thought to facilitate MT formation and organization. A direct connection between RanGTP and MT nucleation could be established via the SAF TPX2, which stimulates branching MT nucleation and thereby generates spindle MTs (1, 9, 10). RanGTP’s role as an upstream regulator of spindle assembly is apparent upon its addition to metaphase-arrested *Xenopus* egg extracts, which leads not only to MT assembly but also the organization of polar MT networks (11–13). Some of these MT networks have been described as “Ran mini spindles”, although their exact architecture shows a large variation across previous studies (2–6). This motivated us to examine how the architecture of Ran-induced MT networks is determined.

While studies in *Xenopus* egg extracts are often conducted in bulk, the influence of extract confinement has mostly been studied with MTs that have been stabilized with Taxol. Taxol addition to bulk *Xenopus* egg extracts induces the formation of MT asters (14), whereas enclosure by droplets of increasing sizes changed the final architecture of MT networks to contractile networks, cortical bundles, vortices, and asters (15, 16). For example, encapsulated taxol-containing extract with a droplet diameter of 100 μm displayed vortices, i.e. there was a strong vortical flow (17). When confined in a rectangular microfluidic channel with a minimum length scale of 100 μm, however, MT asters in the presence of taxol can merge and assemble themselves into a contractile network (18). Independently, physical confinement also affects the organization of MTs with purified components. Encapsulating purified tubulin in the presence of GTP and motors in either microchambers or droplets leads to different MT structures depending on confinement size: at a small confinement size, MTs bent and formed bundles close to the confinement periphery; at a larger confinement size, MTs formed asters by clustering MT ends under the action of motors (16, 19, 20).

All prior studies described above focused on stabilized MTs or MTs that are generated from purified tubulin, and dissected how they interact in the presence of motors and within confinement. Yet, in a cell, MTs are neither stabilized nor generated spontaneously. Instead, they are nucleated from precise locations at the correct cell cycle stage and exhibit dynamic instability during their lifetime. Here, we examine the architecture of MT networks that nucleate dynamic MTs via the RanGTP nucleation pathway within cell-like compartments, generated by encapsulating extract in droplets via microfluidics. We reveal how confinement size and the concentration of the dominant active form of Ran, RanQ69L, regulates the assembly of MT networks. Our work highlights the prominent role of MT nucleation combined with cell-like confinement during the self-organization of MTs, and might have direct implications in nucleation-controlled soft material processing.

## MATERIALS AND METHODS

### *Xenopus* egg extract preparation

Cytoplasmic extracts were prepared from unfertilized *Xenopus* laevis oocytes arrested in metaphase of Meiosis II, as previously described (12, 13) and used within three hours of preparation.

### RanQ69L expression and purification

N-terminal strep-6xHis-TEV mTagBFP2 RanQ69L was cloned into the pST50 vector via Gibson Assembly (New England Biolabs). The plasmid was transformed into *E. coli* Rosetta2 cells (EMD Millipore) for protein expression. Cells were grown until OD600= 0.8 at 37 °C in LB media, and then protein expression was induced using 500 mM IPTG for 18 hours at 16 °C before cells were pelleted. RanQ69L was purified mostly as described (21, 22). Briefly, cells were lysed using an Emulsiflex french press (Avestin) in binding buffer (100 mM Tris-HCl, 450 mM NaCl, 1 mM MgCl_2_, 1 mM EDTA, 2.5 mM PMSF, 6 mM BME, pH 8.75). The lysate was centrifuged at 20,000 x g and the supernatant was loaded onto a StrepTrap HP column (GE Healthcare). Protein was eluted in binding buffer with 2.5 mM D-desthiobiotin, and dialyzed overnight into CSF-XB buffer (10 mM HEPES, 100 mM KCl, 1 mM MgCl_2_, 5 mM EGTA, 10% sucrose w/v, pH 7.7). 200 μM GTP was included in lysis and elution buffers.

### Bulk assay preparation

In a 4 °C cold room, fresh metaphase-arrested egg extracts were supplemented with Alexa-647 labeled tubulin at 0.3 μM final concentration for all the experiments (PurSolutions, VA), and RanQ69L concentration (*[Ran]*) at 5 *μ*M, 10 *μ*M, and 20 *μ*M. The minimum Ran concentration of 5 *μ*M was chosen, because MT nucleation events were rarely observed within a time course of 20 min, consistent with previous work (23).

### Microfluidic channel fabrication and droplet generation

To confine a cell-free extract, we used droplet microfluidics (Fig. 1a). We used standard soft lithographic techniques to fabricate the microchannel in poly(dimethylsiloxane) (PDMS) (24). The microchannel was rendered hydrophobic by treatment with Aquapel (Pittsburgh, PA) to avoid wetting of the wall by droplets. The microchannel consists of an upstream flow-focusing nozzle for generating monodisperse, extract-in-oil droplets (25) (Fig. 1b) and a downstream chamber for collecting the droplets (Fig. 1c). The dispersed phase of the droplets consisted of a mixture of fresh extracts, labelled tubulin, and Ran (see Bulk assay preparation), while the continuous phase was hydrofluoroether HFE-7500 (3M, St. Paul, MN) containing the non-ionic, PEG-based 008-FluoroSurfactant (2% w/w, RanBiotechnologies, USA). The surfactant prevents droplet coalescence by coating the droplet interface with a PEG layer, which is commonly used in droplet-based applications to minimize interactions between proteins and interfaces (20, 26, 27). The flow was driven by a syringe pump (NewEra, NJ). To vary the droplet size, we changed the volumetric flow rates of the dispersed and continuous phases. We characterized the droplet size by their diameter *D*, measured when the droplets were spherical (Fig. 1d and Supplementary Information (SI) Note S1). The dispersity of the generated droplets was < 5% by volume. We fabricated microchannels with various depths *H*, which was always smaller than the droplet diameter *D*. The droplets were thus fully confined by the top and bottom walls of the microchannels and appeared a disc in shape (Fig. 1d). The detailed droplet size *D*, constriction width *W* and depth *H* of the microchannels, and flow rates used for droplet generation are summarized in SI Table S1. For most droplet results (Fig. 1 – Fig. 4), the *H/D* ratio was kept constant at *H/D~0.8*. To test the effect of droplet deformation on the MT networks (Fig. 5), we also tested *H/D~0.4* by reinjecting droplets with *D=110 μm* into a shallow microchannel with *H=40 μm*. All equipment and microchannels were chilled overnight prior to the experiments. We monitored the droplet generation process using a stereoscope in the 4 °C cold room. After sufficient droplets accumulated in the downstream chamber, we stopped the droplet generation immediately by unplugging the inlet tubing. We waited for 10 minutes in the 4 °C cold room until the flow ceased in the microchannel. We then moved the microchannel to a confocal microscope at room temperature (20 °C) for high-resolution imaging. All confocal images were taken at the midplane in the z direction.

**Figure 1.**
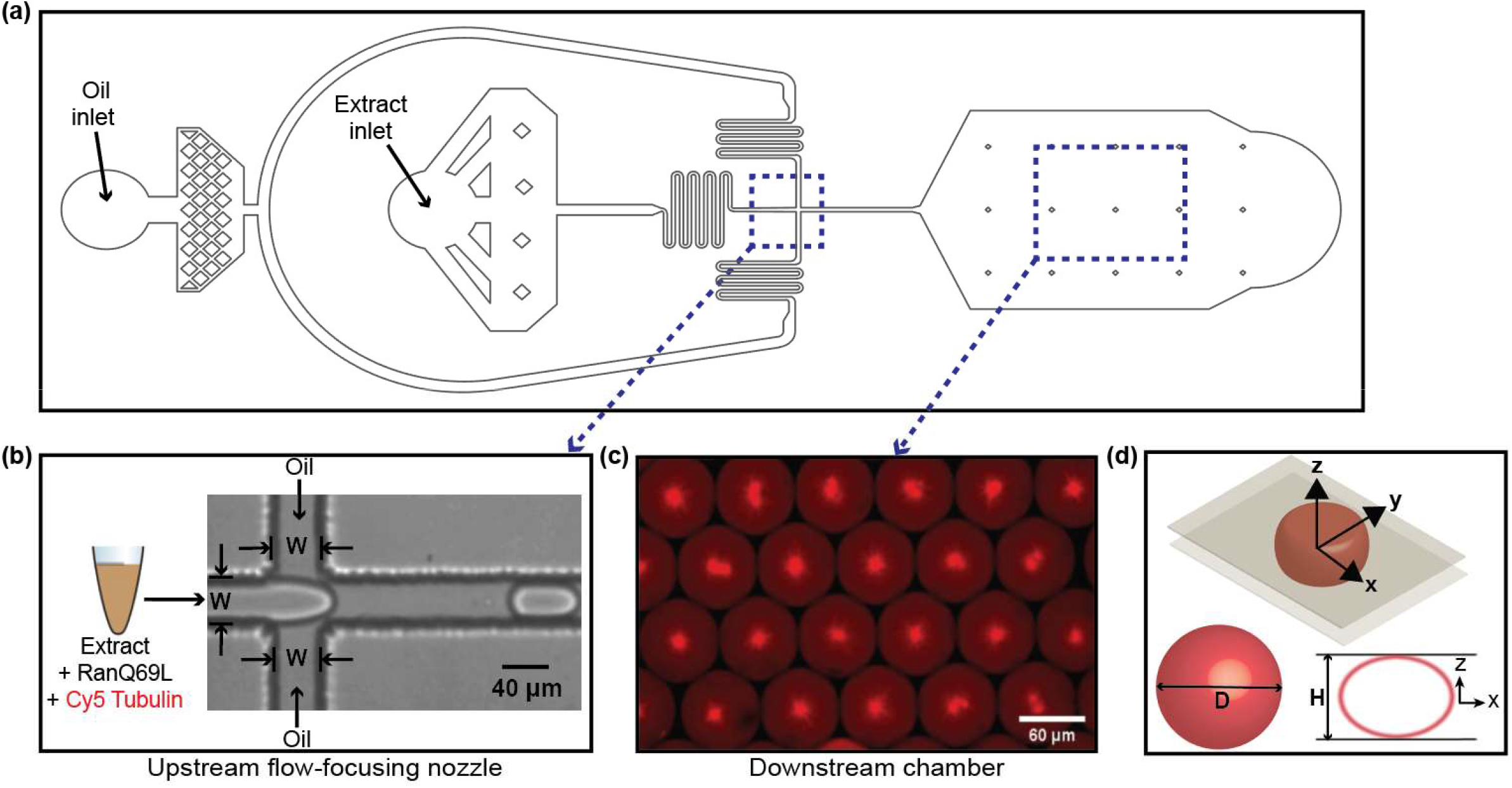
Experimental setup. (a) A snapshot of the microchannel design. The blue dashed boxes in (a) highlight: (b) the upstream flow-focusing channel for generating monodisperse, extract-in-oil droplets; (c) the downstream chamber for collecting and imaging the droplets. (c) is a wide-field confocal image showing the MT network architecture for *D=60 μm* and *[Ran]=20 μM* (same conditions as Fig. 4c) at time *t=14 min*. (d) Schematics of a droplet confined in the microchannel, the definition of droplet diameter *D* measured when the droplet is spherical, and the channel height H.

## RESULTS

### The architecture of MT networks differs drastically when confined in a cell-shaped droplet

To investigate the architecture of RanGTP-induced MT network, we added RanQ69L, loaded with GTP that can no longer be hydrolyzed, to *Xenopus* egg extracts. Formation of MT networks were visualized via the addition of fluorescently labeled tubulin. Within 15 minutes, MTs self-organized into polar networks consisting of interconnected poles and MT bundles (Fig. 2a), as reported previously (3). The overall architecture of RanGTP-induced MT structures remained similar for various Ran concentrations tested (*[Ran]=5 – 20 μM*, Fig. 2a). Between 15 and 40 minutes, the MT network architecture did not vary significantly (SI Fig. S1).

**Figure 2.**
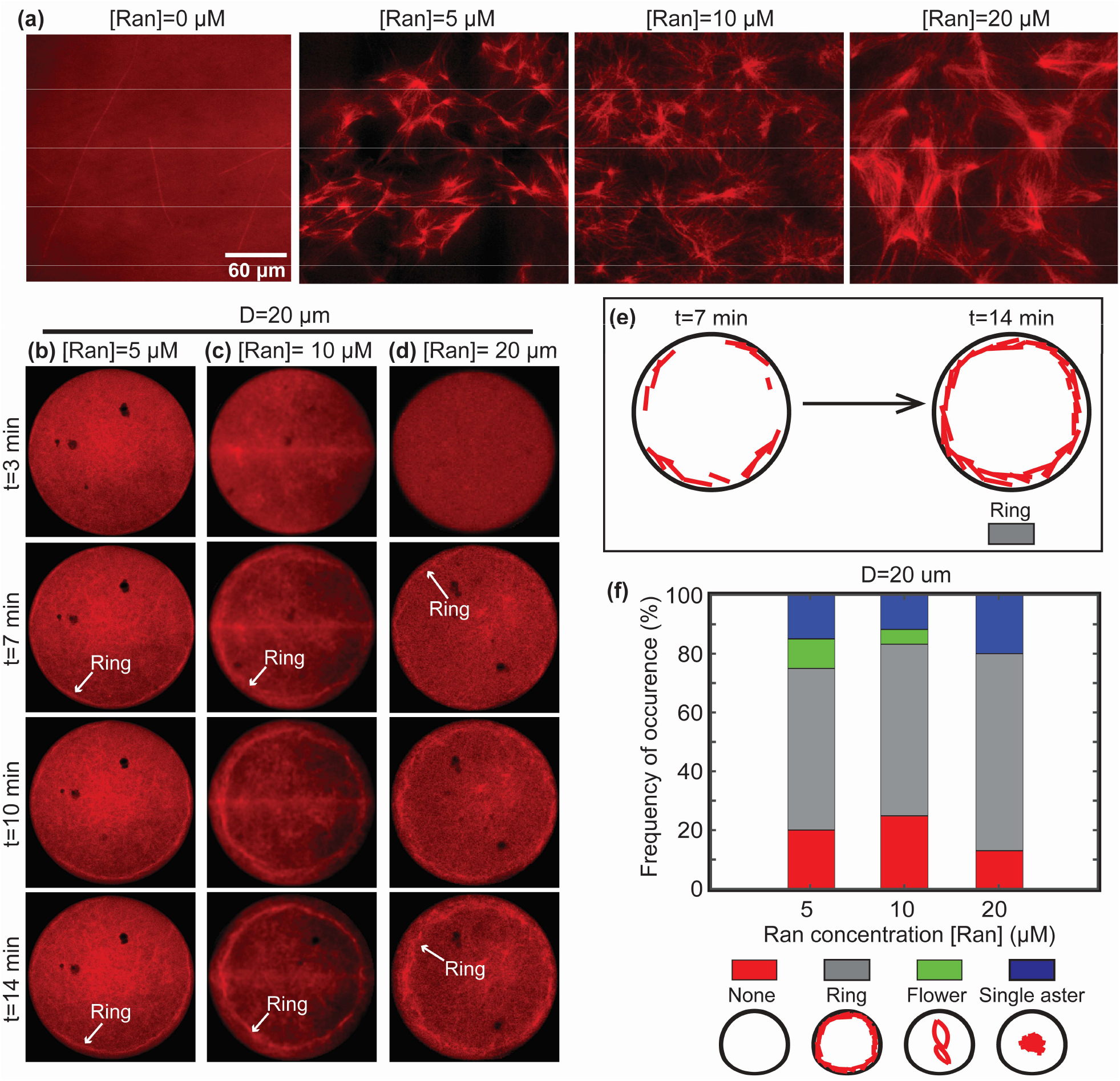
Unconfined MT networks and ring structure at small droplet size. (a) Confocal images showing MT networks (red) in unconfined extracts for Ran concentrations: 0 *μ*M as a control, 5 *μ*M, 10 *μ*M, and 20 *μ*M. The images were taken at time *t=14 min*. The scale bar is associated with (a) only. (b)-(d) The assembly of ring-shaped MT networks in droplets with a diameter of D=20 *μ*m. The Ran concentration *[Ran]* ranges from (b) 5 *μ*M, (c) 10 *μ*M, to (d) 20 *μ*M. (e) A schematic representing the assembly processes of the ring architecture. (f) Frequency of occurrence in percentage of the ring architectures at *[Ran]=5 μ*M, 10 *μ*M, and 20 *μ*M for *D=20 μ*m. Each bar was generated by tracking a total number of ~200 droplets. Each color represents a specific architecture as indicated below.

Next, we examined the assembly of Ran-induced MT networks by time-lapse confocal microscopy in different droplet sizes, generated via microfluidic fabrication (see methods and Fig. 1) and within a physiological range (28, 29). To our surprise, the Ran-induced MT networks differed greatly from the ones formed in bulk extracts when encapsulated in round, finite droplets (Fig. 2b-2-, 3a-3b, and 4a-4d, also see SI Movie 1 - 3). Interestingly, the architecture of primary MT networks formed within 15 minutes and remained overall unchanged. Beyond 15 minutes, we additionally observed the formation of independent, polar MT structures, which were a few orders of magnitude smaller than the primary MT networks and scattered throughout the droplets with *D=110 μm* and *[Ran]=10 μM*) (SI Fig. S2). Upon varying the droplet diameter and Ran concentration, most changes occurred in the larger, primary MT structures within the first 15 minutes, as reported below.

**Figure 3.**
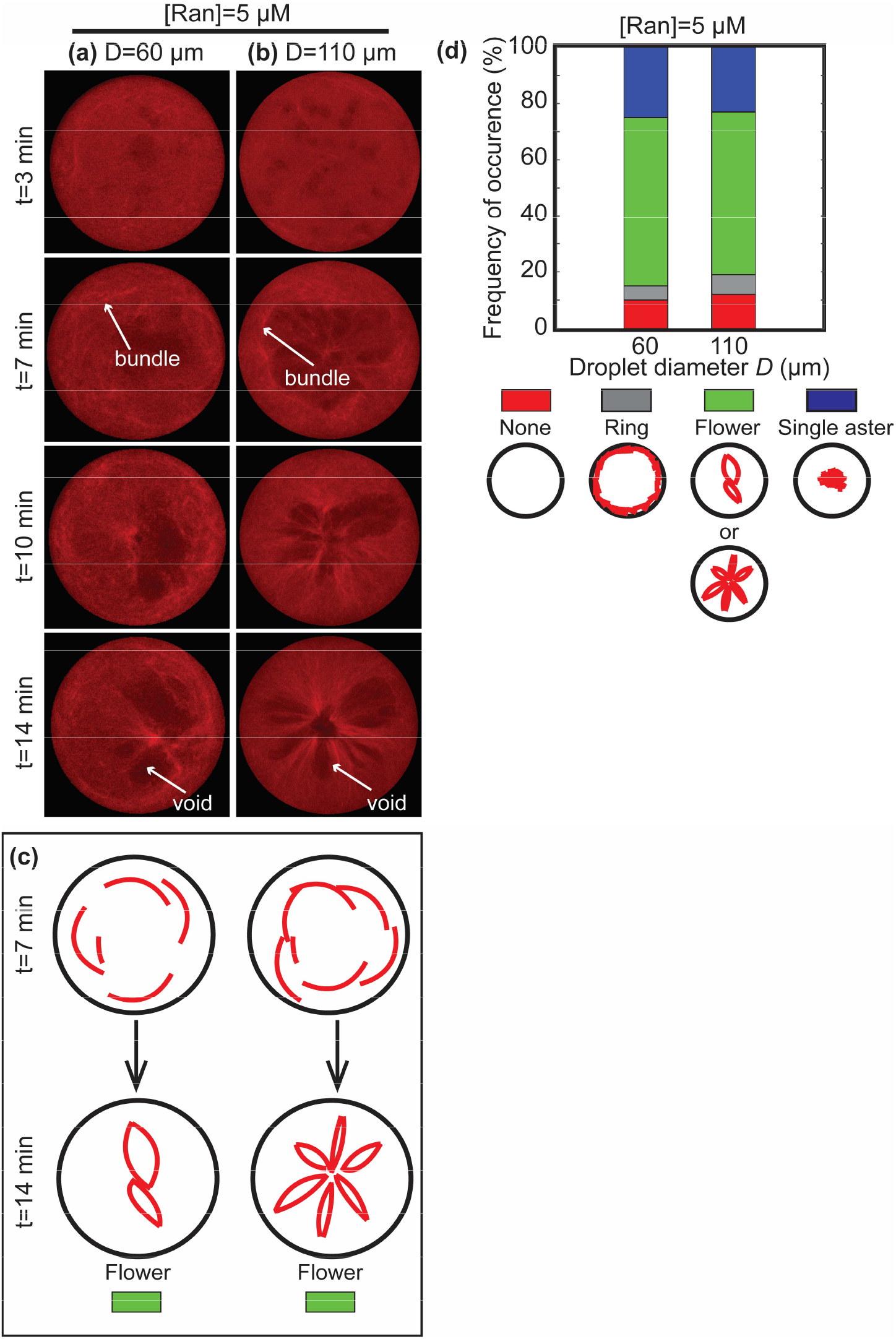
Flower structure at *[Ran]=5 μM* for D= 60 *μ*m and *D=110 μ*m. (a)-(b) Confocal images showing the assembly of flower-shaped MT networks inside droplets. The Ran concentration is 5 *μ*M and the droplet diameters are (a) D=60 *μ*m and (b) D=110 *μ*m, respectively. (c) Schematics showing the assembly of the flower-shaped MT networks. (d) Frequency of occurrence in percentage of the flower architecture at *[Ran]=5 μ*M for D=60 *μ*m and (b) D=110 *μ*m, respectively. Each color represents a specific MT network architecture below.

### Small droplet size (*D=20 μm*) leads to a ring-shaped architecture of MT networks

At a droplet diameter *D=20 μm*, MTs assembled into a ring near the droplet boundary (Fig. 2b-2d). Starting at *t=7 min*, a ring of MTs became visible near the droplet boundary and maintained its architecture thereafter. This is consistent with a previous study, in which Taxol-stabilized MTs formed a cortical array close to the boundary, which was caused by MTs bending along the boundary when the length of the MTs became comparable to the dimension of confining space (15, 16). For droplets with a diameter of *D=20 μm*, the ring-shaped architecture is insensitive to changes in Ran concentration (Fig. 2b – 2d), and robust over dozens of droplets (SI Fig. S3). Thus, the formation of the ring-shaped architecture is largely determined by confinement size rather than the nucleation activity of MTs for small droplet diameters. At higher Ran concentrations of 10 and 20 uM, additional internal MT structures start to appear over time (Fig. 2f and SI Fig. S3). Importantly, the MT architecture in confined small droplets differs greatly from the same sample in an unconfined sample chamber.

### Droplet size affects MT network architecture at low Ran concentrations

During spindle assembly, a gradient of Ran concentration exists in the vicinity of the chromosomes (1, 8). Based on this, we hypothesized that, in addition to the droplet size, the variation of Ran concentration regulates the architecture of MT networks for larger droplet sizes. To test our hypothesis, we varied the Ran concentration in droplets with *D=60 μm* and *D=110 μm*.

At the low Ran concentration of 5 *μM* and in droplets with the diameters of 60 and 110 *μm* (Fig. 3a – 3b; also see SI Movie S2), MTs assembled into a network partitioned by “void areas” (white arrows in Fig. 3a – 3b at *t=14 min*), which exhibited a low MT density and precluded the assembly of MTs. As a result, the architecture of the network resembled the shape of a flower. The number of partitions or void areas scaled with the droplet size and was robust over dozens of droplets (see SI Fig. S4 – S5 and SI Movie S4). On average, droplets with *D=60 μm* have two or three partitions, and the larger droplets with *D=110 μm* have five to seven partitions. The assembly of the flower-shaped architecture was achieved through contraction of the MT network. From *t=3 min* to *t=7 min*, most MTs nucleated close to the boundary of the droplets, rather than homogeneously inside the droplets, and formed bundled MTs (white arrows in Fig. 3a – 3b at *t=7 min*). This resulted in a brighter region with a higher MT density that surrounds a darker region with a lower MT density. Subsequently, the perimeter of MTs contracted in an asymmetric way, such that the MT network became partitioned by the darker void areas into a flower-shaped architecture.

### Ran concentration affects MT network architecture at medium and large droplet sizes

When increasing the Ran concentration to 10 *μ*M and 20 *μ*M within droplet diameters of 60 and 110 μm, MTs primarily assembled into asters, i.e., a radial array of MTs that is centered in the droplet. The detailed characteristics of the assembly process, however, depended on both droplet diameter *D* and Ran concentration. To illustrate the assembly process, we focus on comparing the morphologies of MT networks formed at two time points: an intermediate state at *t= 7 min* characterized by ‘clusters’ of MTs that are spatially separated, and a steady state at *t=14 min* with a characteristic aster of MTs (Fig. 4a – 4d). We define ‘clusters’ as the astral MT networks’ subsets that are still undergoing transient spatial movements and morphological changes *t=7 min*, leading eventually to the steadystate MT asters at *t=14 min*.

**Figure 4.**
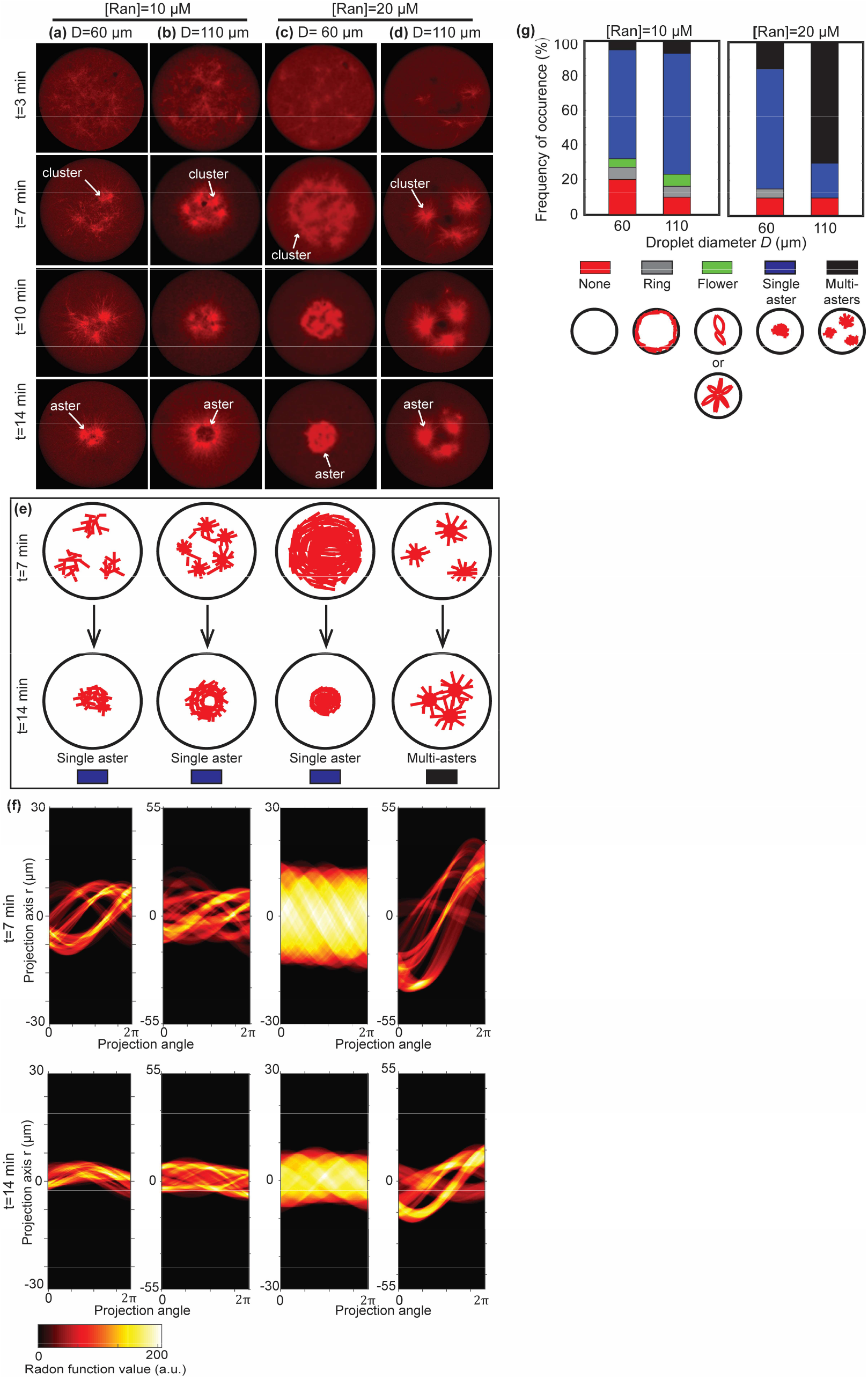
Aster MT networks. (a)-(d) Time-lapse confocal images showing the assembly of asters inside droplets. (e) Schematics showing the assembly of the asters. (f) Radon transform of the architectures at *t=7 min* (top row) and *t=14 min* (bottom row), respectively. (g) Frequency of occurrence of the aster architecture. Each color represents a specific MT network architecture shown below.

At a Ran concentration of 10 μM and in droplets with diameters of 60 μm and 110 μm (Fig. 4a – 4b; also see SI Movie S3), MTs first bundled into a few spatially-separate clusters at *t=7 min*. They subsequently aggregated to the center of droplets and formed radial asters at *t=14 min*.

At a Ran concentration of 20 *μM* and a droplet diameter of 60 *μ*m (Fig. 4c), in contrast to the relatively separate clusters (Fig. 4a – 4b at *t=7 min*) at *[Ran]=10 μM*, MTs formed an increased number of clusters packing densely and appearing to be interconnected at *t=7 min*, likely due to enhanced MT nucleation activities at a higher Ran concentration. The difference between the separate and interconnected clusters at *t=7 min* is further supported by the Radon transform of the images (Fig. 4f, top row), where the separated clusters show sinusoidal wave patterns, and the interconnected clusters show a homogeneous stripe. With time, the entire network of interconnected clusters contracted radially, leading to a single aster at *t=14 min*.

Further increasing the droplet diameter to 110 *μm* at *[Ran]=20 μM* (Fig. 4d), we observed that the clusters again became separate at *t=7 min* (Fig. 4d), supported by the sinusoidal wave in their corresponding Radon transform at *t=7 min* (Fig 4g). Subsequently, the separate clusters moved towards the center of the droplet. Rather than forming a single aster as in the previous cases (Fig. 4a – 4c at *t=14 min*), they did not collapse, giving rise to a network of multiple asters coexisting inside the droplet. The contrast between the single-aster droplets versus the multi-aster droplets can also be seen by comparing their Radon transforms at *t=14 min* (Fig. 4f, bottom row), where the multi-aster case shows a clear sinusoidal wave in comparison to the single aster cases. The assembly of asters inside droplets is robust over dozens of droplets (Fig. 4g, SI Fig. S6; also see SI Movie S5).

### Effect of motor activity and droplet deformation

To test whether motor activity is required for the assembly of MT networks in our system, we inhibited motor activities by adding vanadate (Fig. 5a and SI Movie S6) (22). MTs formed, but failed to assemble into a larger network with a defined architecture inside the droplets. Instead, we observed scattered MT bundles, which are expected to form from branching MT nucleation that gives rise to characteristic branched MT networks (22). Therefore, within a sufficiently large confinement for MTs to grow freely without boundary compression, the activities of motors are necessary to pull separate nucleation centers together to assemble Ran-induced MT networks.

**Figure 5.**
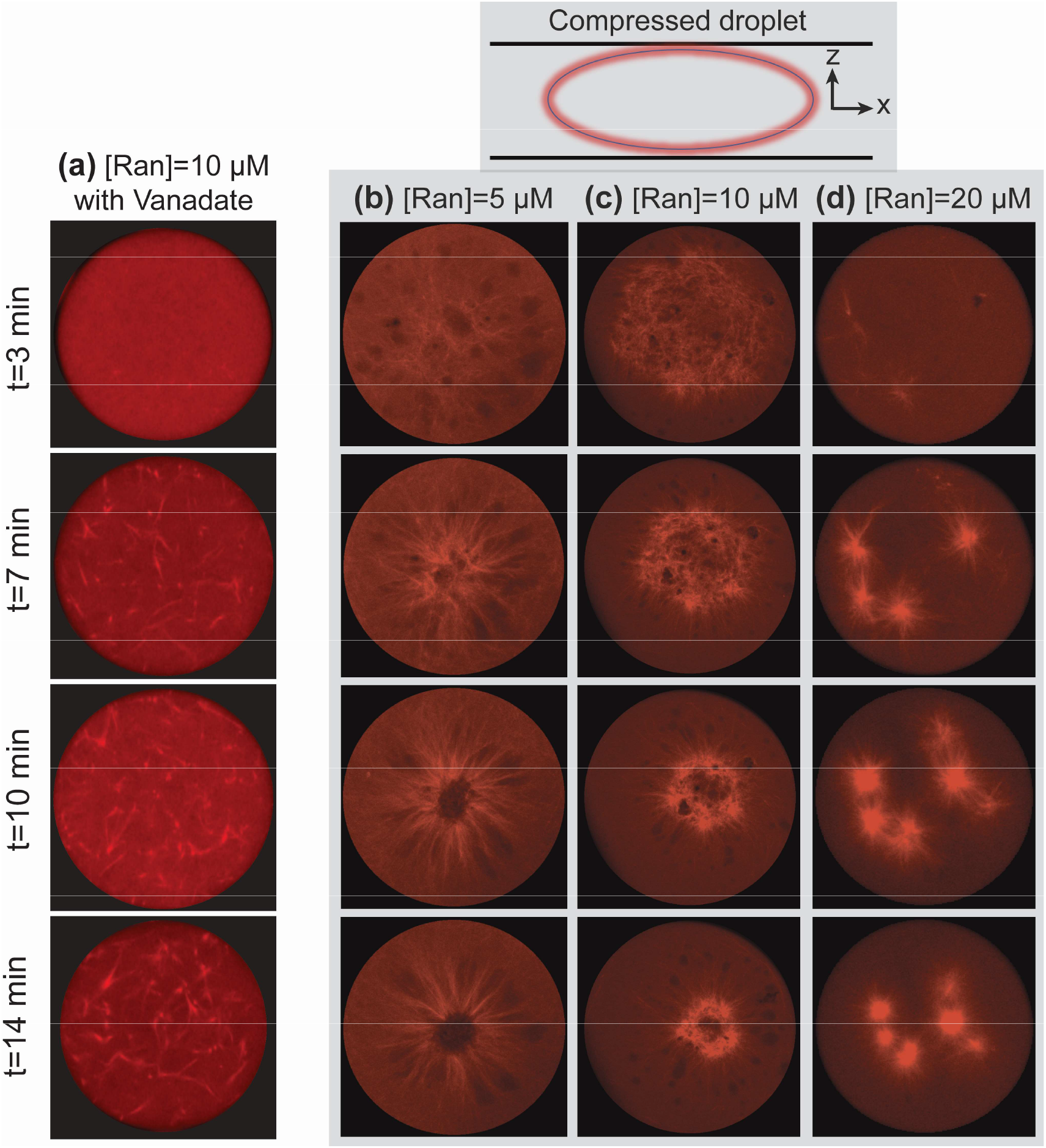
Effect of droplet deformation and inhibiting motor activities. (a) Effect of inhibiting motor proteins in confined extracts. The droplets have a diameter of 110 *μm*. The concentration of Vanadate in the droplets is 500 nM. See SI Movie S6 for more details. (b)-(d) Time-lapse confocal images showing the assembly of MT networks in compressed droplets at three Ran concentrations: (b) 5 *μ*M, (c) 10 *μ*M, and (d) 20 *μ*M. For the compressed droplet, *H/D~0.4*. The grey background highlights the compressed-droplet cases. The size of the compressed droplets in (b)-(d) is the same as an uncompressed droplet with *D=110 μm*.

To understand how the shape of droplets affect the MT network assembly, we deformed the droplets without changing their volume by reinjecting the droplets into a chamber with a small depth, such that the channel depth-to-droplet-diameter ratio became *H/D~0.4* (see Materials and Methods). The droplets deformed into a pancake shape while maintaining their volume. For the same droplet size and Ran concentration, we observe that the assembly processes and the steady-state architectures of the MT networks in the compressed droplets (Fig. 5b – d) are consistent with those in the uncompressed droplets (Fig. 3b, Fig. 4b, and Fig. 4d). These observation are in agreement with a prior study (30) that reported confinement volume, not confinement shape, drives spindle scaling and limited cytoplasmic components affect the MT network assembly.

## DISCUSSION

Here, we found that MT architecture depends on cytoplasmic volume, MT nucleation, and motor activity. Specifically, all MT architectures differ from the ones formed in bulk extract. Thus, we summarize the steady-state (*t=14 min*) architecture of Ran-induced MT networks as a function of droplet size *D* and Ran concentration *[Ran]* in a regime map (Fig. 6). We have identified four types of architectures: rings, flowers, single asters, and multiple asters. The assembly of the architectures is robust across many droplets with the same droplet size and Ran concentration.

**Figure 6.**
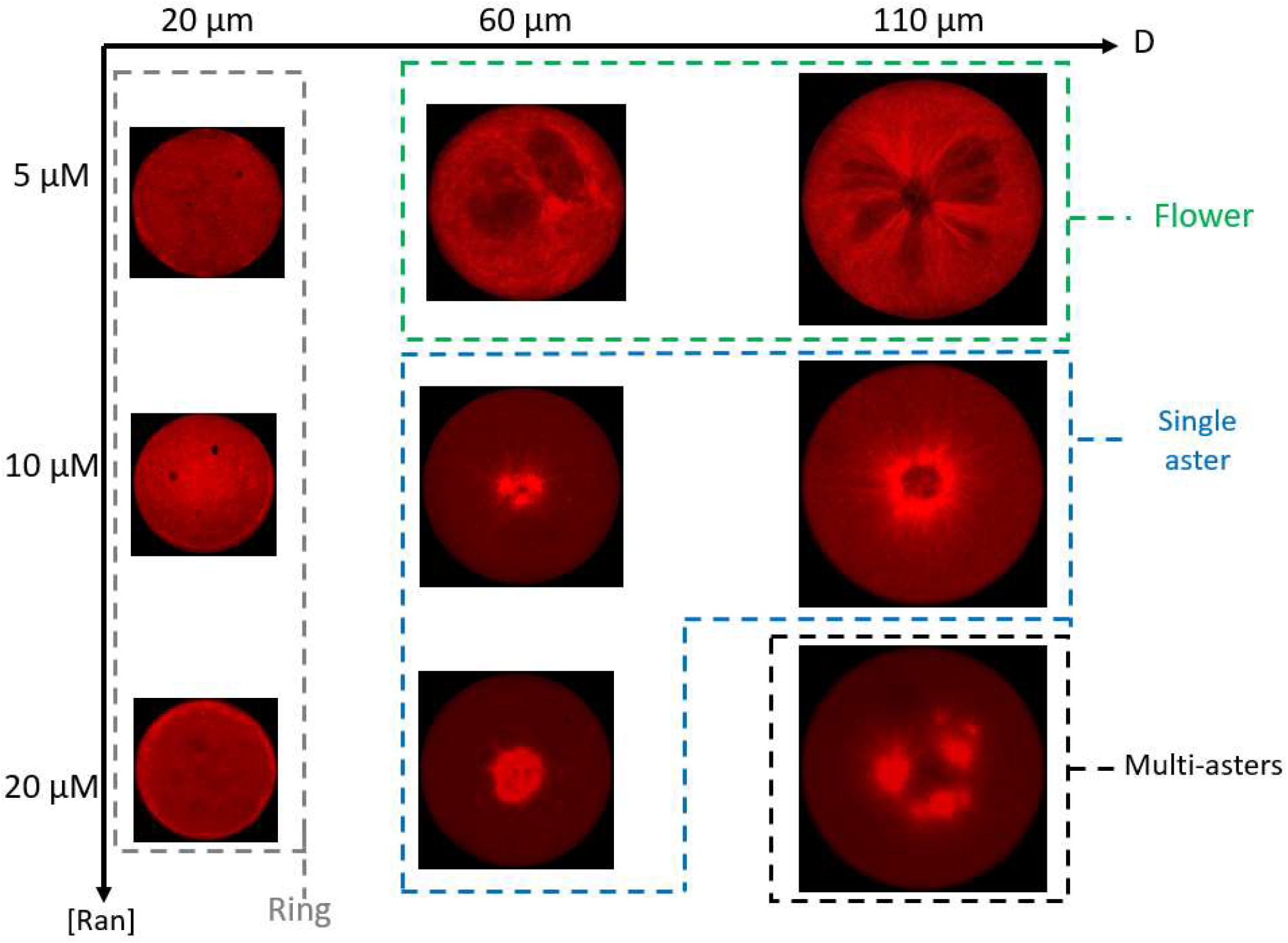
Regime maps summarizing our results. Regime maps summarizing the assembly of MT networks at steady state *t=14 min*. The horizontal axis denotes the droplet diameter, and the vertical axis denotes the Ran concentration. The colored dashed boxes highlight different MT network architectures.

### From rings to asters as *D* increases: pure confinement effect

The formation of the ring architecture may largely be due to a high degree of confinement imposed by the small droplet size. At *D=20 μm*, the length scale of the confining space is close to the reported length scale of Ran-induced MTs during spindle assembly (31, 32). When the MTs grow within the small droplet, their ends eventually touch the droplet boundary. As the growth continues, the MTs buckle and bend along the droplet boundary under the compressive force exerted by the droplet interfacial tension, leading to the formation of the ring structure. A similar ring-assembly process has also been observed in prior studies with confined, stabilized MTs (15, 16).

At *D=60 μm* and *D=110 μm*, more space is available for MT growth and motor-mediated transport of MTs. When the confining space is sufficiently large (*D=60 μm* or *D=110 μm*), molecular motors organize MTs through MT bundling and directed MT transport along MTs (15, 16, 19, 33). Detailed mechanisms of motor activities are beyond the scope of our study and reported in other studies (1, 19, 33–35). In order to merge branched MT networks from a scattered distribution, motor activity is required to organize smaller bundles into larger-scale MT networks, whose architectures range from flowers, to asters, to multipolar arrays.

An architecture that has not been observed before are the flower-like networks. The void areas could potentially be formed by lipids. What exactly are these shapes and whether they occur in a cell is an interesting future area of research.

### MT nucleation with limited resources can lead to different MT architectures

A possible explanation for the variation from flowers, to asters, to multipolar arrays at larger droplet sizes and increasing Ran concentrations can be the competition between MT nucleation and limited resources. The higher the Ran concentration, the more MT nucleation is induced, while the limited resources are a consequence of confinement. Prior studies have suggested that the limited tubulin of confined extracts are the main determinant of the size scaling of mitotic spindles (30, 36). We thus focus our discussion of limited resources on tubulin, the basic structural unit of microtubules.

At a low Ran concentration, there are fewer MT nucleation events. Each MT has sufficient tubulin that guarantees its maximum growth, and thus MTs have a long average length favoring the formation of long, bundled filaments, consistent with our observation at *[Ran]=5 μM* (Fig. 3a – 3b). As the Ran concentration increases, a higher rate of MT nucleation leads to a larger number of MTs during the same amount of time, possibly making the tubulin available in confined extracts limiting for full growth of all nucleated MTs. Consequently, the average length of the MTs becomes shorter than at a lower Ran concentration. In these conditions, motors pack the MTs efficiently into a single aster (15, 16, 33, 37, 38), consistent with our observation at *[Ran]=10 μM* (Fig. 4a – 4b). When the Ran concentration was further increased to *20* μM, MTs are organized into multiple asters coexisting in a droplet (Fig. 4d). which could be due to steric interactions among MTs that prevent excess MTs from packing into a single aster.

The hypothesis of limited resources is supported by the experiments in which droplets were deformed (Fig. 5b – 5d). Given the same droplet volume and Ran concentration, the MT networks exhibited almost identical architectures, despite having a more ovoid shape. Similar observations were made in a prior study (30), where confinement volume, not confinement shape, drives the spindle scaling, a result of the limiting pool of cytoplasmic components.

Our results highlight the critical role of confinement in regulating MT network assembly. Most importantly, we show that MT nucleation is a major determinant of the resulting MT architecture in confined space. This has not been taken into account when working with stabilized MTs and motors. Our work suggests that more studies are needed to further understand how MT nucleation from defined locations and at precise times in a cell leads to a functional MT architecture. Last, our work illustrates a rather simple strategy to tune the architecture of MT networks by changing the droplet size and Ran concentration.

## Author Contributions

Y.G., S.P., and H.A.S. conceived the project, designed the experiments, and wrote the manuscript. S.P. and H.A.S. supervised and guided the research. Y.G. fabricated the devices, prepared the samples, performed the experiments, and analyzed the data. S.S. and B.C. helped with reagents and support. All authors contributed to the discussions and interpretation of the experimental results, and edited the manuscript.

## Acknowledgements

We are grateful to all members of the Petry Lab and the Stone Lab for insightful discussions. This work was supported by the Princeton Catalysis Initiative (to H.A. S. and S.P.), the NIH New Innovator Award, and the David and Lucile Packard Foundation (all to S.P.).

## Notes

### Competing Interest Statement

The authors have declared no competing interest.

## Reference

1. Petry, S. 2016. Mechanisms of mitotic spindle assembly. Annu. Rev. Biochem. 85:659–683.

2. Zhang, C., M. Hughes, and P. R. Clarke. 1999. Ran-GTP stabilises microtubule asters and inhibits nuclear assembly in Xenopus egg extracts. J. Cell Sci. 112(14):2453–2461.

3. Carazo-Salas, R. E., G. Guarguaglini, O. J. Gruss, A. Segref, E. Karsenti, and I. W. Mattaj. 1999. Generation of GTP-bound Ran by RCC1 is required for chromatin-induced mitotic spindle formation. Nature 400(6740):178.

4. Ohba, T., M. Nakamura, H. Nishitani, and T. Nishimoto. 1999. Self-organization of microtubule asters induced in Xenopus egg extracts by GTP-bound Ran. Science 284(5418):1356–1358.

5. Wilde, A., and Y. Zheng. 1999. Stimulation of microtubule aster formation and spindle assembly by the small GTPase Ran. Science 284(5418):1359–1362.

6. Carazo-Salas, R. E., O. J. Gruss, I. W. Mattaj, and E. Karsenti. 2001. Ran–GTP coordinates regulation of microtubule nucleation and dynamics during mitotic-spindle assembly. Nature Cell Biol. 3(3):228.

7. Desai, A., and T. J. Mitchison. 1997. Microtubule polymerization dynamics. Annu. Rev. Cell Dev. Biol. 13(1):83–117.

8. Kalab, P., K. Weis, and R. Heald. 2002. Visualization of a Ran-GTP gradient in interphase and mitotic Xenopus egg extracts. Science 295(5564):2452–2456.

9. Alfaro-Aco, R., A. Thawani, and S. Petry. 2017. Structural analysis of the role of TPX2 in branching microtubule nucleation. J. Cell Biol. 216(4):983–997.

10. Gruss, O. J., R. E. Carazo-Salas, C. A. Schatz, G. Guarguaglini, J. Kast, M. Wilm, N. Le Bot, I. Vernos, E. Karsenti, and I. W. Mattaj. 2001. Ran induces spindle assembly by reversing the inhibitory effect of importin α on TPX2 activity. Cell 104(1):83–93.

11. Desai, A., A. Murray, T. J. Mitchison, and C. E. Walczak. 1998. The use of Xenopus egg extracts to study mitotic spindle assembly and function in vitro. Methods in cell biology. Elsevier, pp. 385–412.

12. Hannak, E., and R. Heald. 2006. Investigating mitotic spindle assembly and function in vitro using Xenopus laevis egg extracts. Nat. Protoc. 1(5):2305.

13. Good, M. C. 2016. Encapsulation of Xenopus egg and embryo extract spindle assembly reactions in synthetic cell-like compartments with tunable size. The Mitotic Spindle. Springer, pp. 87–108.

14. Verde, F., J.-M. Berrez, C. Antony, and E. Karsenti. 1991. Taxol-induced microtubule asters in mitotic extracts of Xenopus eggs: requirement for phosphorylated factors and cytoplasmic dynein. J. Cell Biol. 112(6):1177–1187.

15. Pinot, M., F. Chesnel, J. Kubiak, I. Arnal, F. Nedelec, and Z. Gueroui. 2009. Effects of confinement on the self-organization of microtubules and motors. Curr. Biol. 19(11):954–960.

16. Baumann, H., and T. Surrey. 2014. Motor-mediated cortical versus astral microtubule organization in lipid-monolayered droplets. J. Biol. Chem. 289(32):22524–22535.

17. Suzuki, K., M. Miyazaki, J. Takagi, T. Itabashi, and S. i. Ishiwata. 2017. Spatial confinement of active microtubule networks induces large-scale rotational cytoplasmic flow. Proc. Natl. Acad. Sci. USA 114(11):2922–2927.

18. Foster, P. J., S. Fürthauer, M. J. Shelley, and D. J. Needleman. 2015. Active contraction of microtubule networks. eLife 4:e10837.

19. Nedelec, F., T. Surrey, A. C. Maggs, and S. Leibler. 1997. Self-organization of microtubules and motors. Nature 389(6648):305.

20. Juniper, M. P., M. Weiss, I. Platzman, J. P Spatz, and T. Surrey. 2018. Spherical network contraction forms microtubule asters in confinement. Soft matter 14(6):901–909.

21. Weis, K., C. Dingwall, and A. I. Lamond. 1996. Characterization of the nuclear protein import mechanism using Ran mutants with altered nucleotide binding specificities. EMBO J. 15(24):7120–7128.

22. Petry, S., A. C. Groen, K. Ishihara, T. J. Mitchison, and R. D. Vale. 2013. Branching microtubule nucleation in Xenopus egg extracts mediated by augmin and TPX2. Cell 152(4):768–777.

23. Hoffmann, C., E. Mazari, S. Lallet, R. Le Borgne, V. Marchi, C. Gosse, and Z. Gueroui. 2013. Spatiotemporal control of microtubule nucleation and assembly using magnetic nanoparticles. Nat. Nanotechnol. 8(3):199.

24. Tang, S. K., and G. M. Whitesides. 2010. Basic microfluidic and soft lithographic techniques. McGraw-Hill.

25. Anna, S. L., N. Bontoux, and H. A. Stone. 2003. Formation of dispersions using “flow focusing” in microchannels. Appl. Phys. Lett. 82(3):364–366.

26. Oakey, J., and J. C. Gatlin. 2018. Microfluidic Encapsulation of Demembranated Sperm Nuclei in Xenopus Egg Extracts. Cold Spring Harb. Protoc. 2018(8):pdb. prot102913.

27. Guan, Y., Z. Li, S. Wang, P. M. Barnes, X. Liu, H. Xu, M. Jin, A. P. Liu, and Q. Yang. 2018. A robust and tunable mitotic oscillator in artificial cells. eLife 7:e33549.

28. Wühr, M., Y. Chen, S. Dumont, A. C. Groen, D. J. Needleman, A. Salic, and T. J. Mitchison. 2008. Evidence for an upper limit to mitotic spindle length. Curr. Biol. 18(16):1256–1261.

29. Montorzi, M., M. H. Burgos, and K. H. Falchuk. 2000. Xenopus laevis embryo development: Arrest of epidermal cell differentiation by the chelating agent 1, 10 - phenanthroline. Mol. Reprod. Dev. 55(1):75–82.

30. Hazel, J., K. Krutkramelis, P. Mooney, M. Tomschik, K. Gerow, J. Oakey, and J. Gatlin. 2013. Changes in cytoplasmic volume are sufficient to drive spindle scaling. Science 342(6160):853–856.

31. Brugués, J., V. Nuzzo, E. Mazur, and D. J. Needleman. 2012. Nucleation and transport organize microtubules in metaphase spindles. Cell 149(3):554–564.

32. Decker, F., D. Oriola, B. Dalton, and J. Brugués. 2018. Autocatalytic microtubule nucleation determines the size and mass of Xenopus laevis egg extract spindles. eLife 7:e31149.

33. Surrey, T., F. Nédélec, S. Leibler, and E. Karsenti. 2001. Physical properties determining self-organization of motors and microtubules. Science 292(5519):1167–1171.

34. Roostalu, J., J. Rickman, C. Thomas, F. Nédélec, and T. Surrey. 2018. Determinants of polar versus nematic organization in networks of dynamic microtubules and mitotic motors. Cell 175(3):796–808. e714.

35. Wittmann, T., A. Hyman, and A. Desai. 2001. The spindle: a dynamic assembly of microtubules and motors. Nature Cell Biol. 3(1):E28–E34.

36. Good, M. C., M. D. Vahey, A. Skandarajah, D. A. Fletcher, and R. Heald. 2013. Cytoplasmic volume modulates spindle size during embryogenesis. Science 342(6160):856–860.

37. Heald, R., R. Tournebize, T. Blank, R. Sandaltzopoulos, P. Becker, A. Hyman, and E. Karsenti. 1996. Self-organization of microtubules into bipolar spindles around artificial chromosomes in Xenopus egg extracts. Nature 382(6590):420.

38. Cai, S., L. N. Weaver, S. C. Ems-McClung, and C. E. Walczak. 2009. Kinesin-14 family proteins HSET/XCTK2 control spindle length by cross-linking and sliding microtubules. Mol. Biol. Cell 20(5):1348–1359.

