## Supplementary Information for "Confinement size determines the architecture of Ran-induced microtubule networks"

**Note S1 Geometrical parameters of the microchannels and applied flow rates**

For the results presented in our study, the droplet size is characterized by *D,* the nominal diameter of the droplet when it is spherical. Since *D* is smaller than *H*, the microchannel depth, the droplet is compressed into a pancake shape by the upper and lower walls of the microchannel (Note S1 Fig. 1d). The deformation of a droplet confined between two parallel plates has been studied in detail in prior works. These works suggest the relationship between *D* and *D'*, the diameter of the droplet periphery at its midplane can be described by Eq. S1.

$$\left(\frac{D}{H}\right)^2\left(q+\frac{\pi}{4}\sqrt{2q}+1-\frac{3\pi^2}{32}\right)=\frac{D'}{D}$$

$$\text{where } q=\frac{1}{3}\left(\frac{H}{D}\right)^3-\frac{1}{3}+\frac{\pi^2}{32} \qquad \text{Eq. (S1)}$$

To measure the droplet size *D*, we first measured *D'* from confocal images and use Eq. S1 to convert *D'* into *D*.

**Table *S*1. Summary of the flow rates, microchannel geometrical parameters, and droplet size. $Q_d$ and $Q_c$ denote the disperse and continuous phase flow rate, respectively.**

| *D (μm)* | *W (μm)* | *H (μm)* | $Q_d$ *(mL/hr)* | $Q_c$ *(mL/hr)* |
|---|---|---|---|---|
| 20 | 20 | 15 | 0.3 | 1 |
| 60 | 40 | 50 | 0.3 | 0.3 |
| 110 | 90 | 90 | 0.3 | 0.4 |

**Figure S1. Ran-induced MT networks in bulk extracts at *t=40 min***

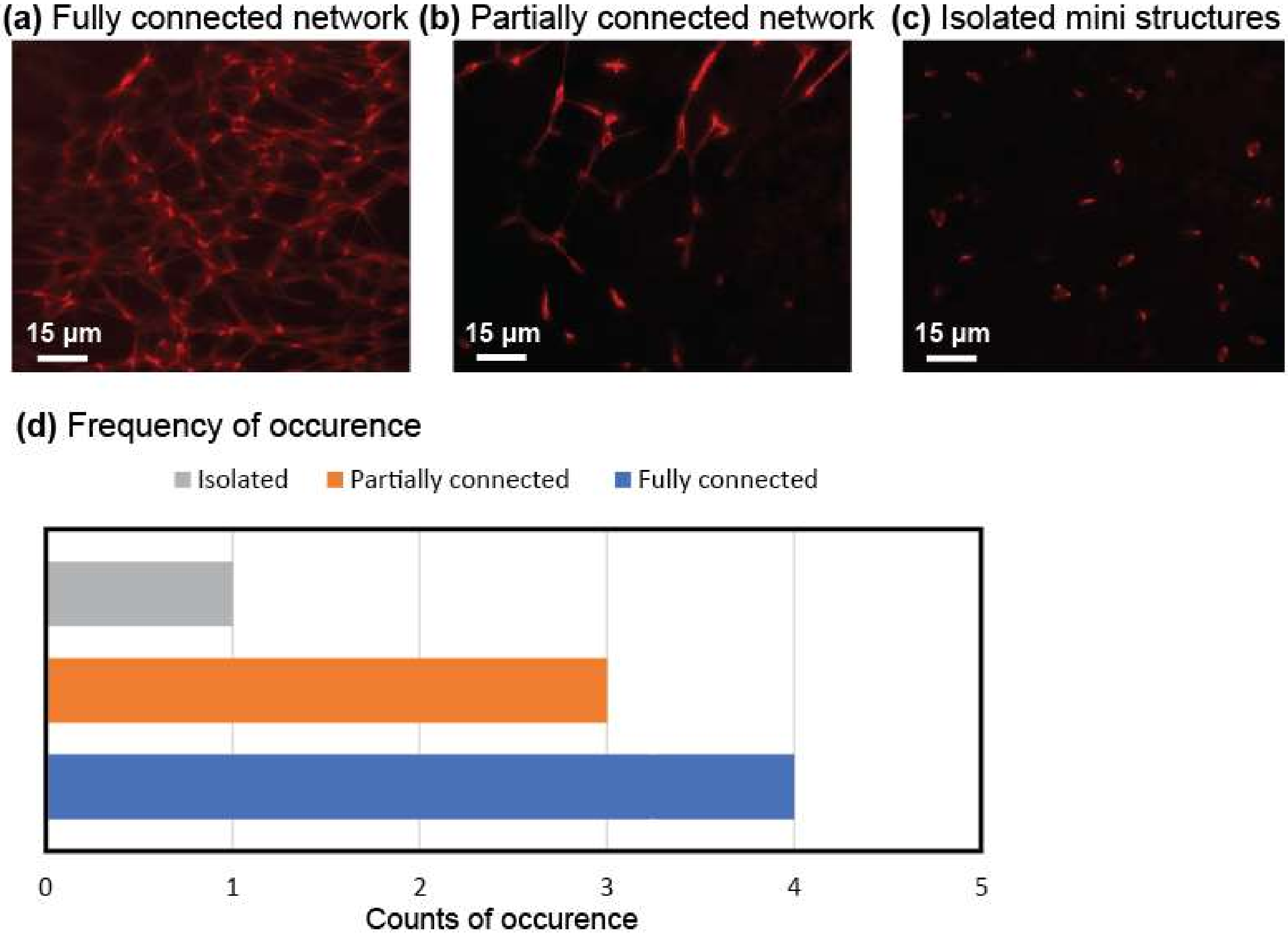


Figure S1. 3D projection of z-scan confocal images taken at *t=40 min* for unconfined extracts. The Ran concentration is *[Ran]=10 μM*. Within the field of view, three types of MT network architectures can be observed: (a) A fully connected network, where the MTs were assembled into dense poles interconnected by thick MT bundles; (b) A partially connected network, where isolated structures and interconnected networks coexisted; (c) Completely isolated mini structures. The frequency of occurrence (based on 8 experiments, each corresponds to one preparation of extract) of the architectures is summarized in (d). Most of our experiments show either fully or partially connected networks.

**Figure S2. Ran-induced MT networks in droplets at *t=40 min***

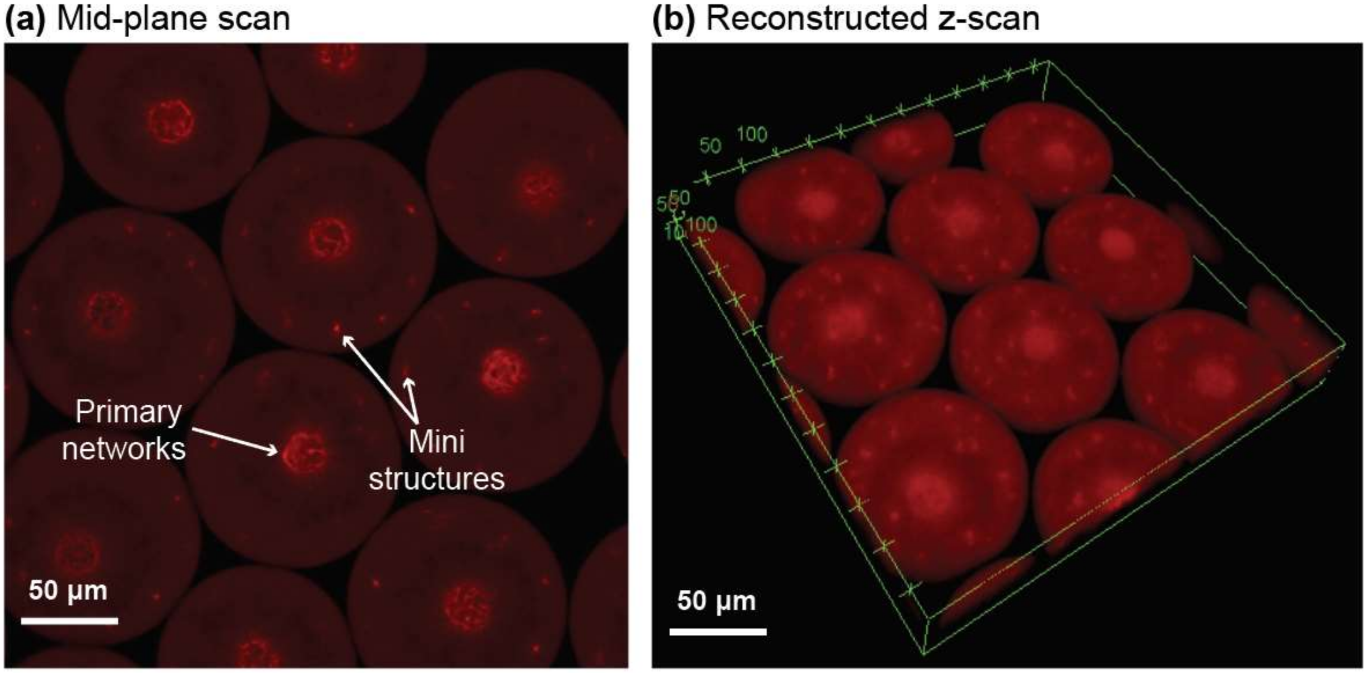


Confocal images taken at *t=40 min* for encapsulated extracts. The droplet size is $D=110\ \mu m$, and the ran concentration is $[Ran]=10\ \mu M$. (a) The image was taken at the mid plane. (b) Reconstructed 3D images from z-scan.

**Figure S3. Wide-field images showing the assembly of MT rings**

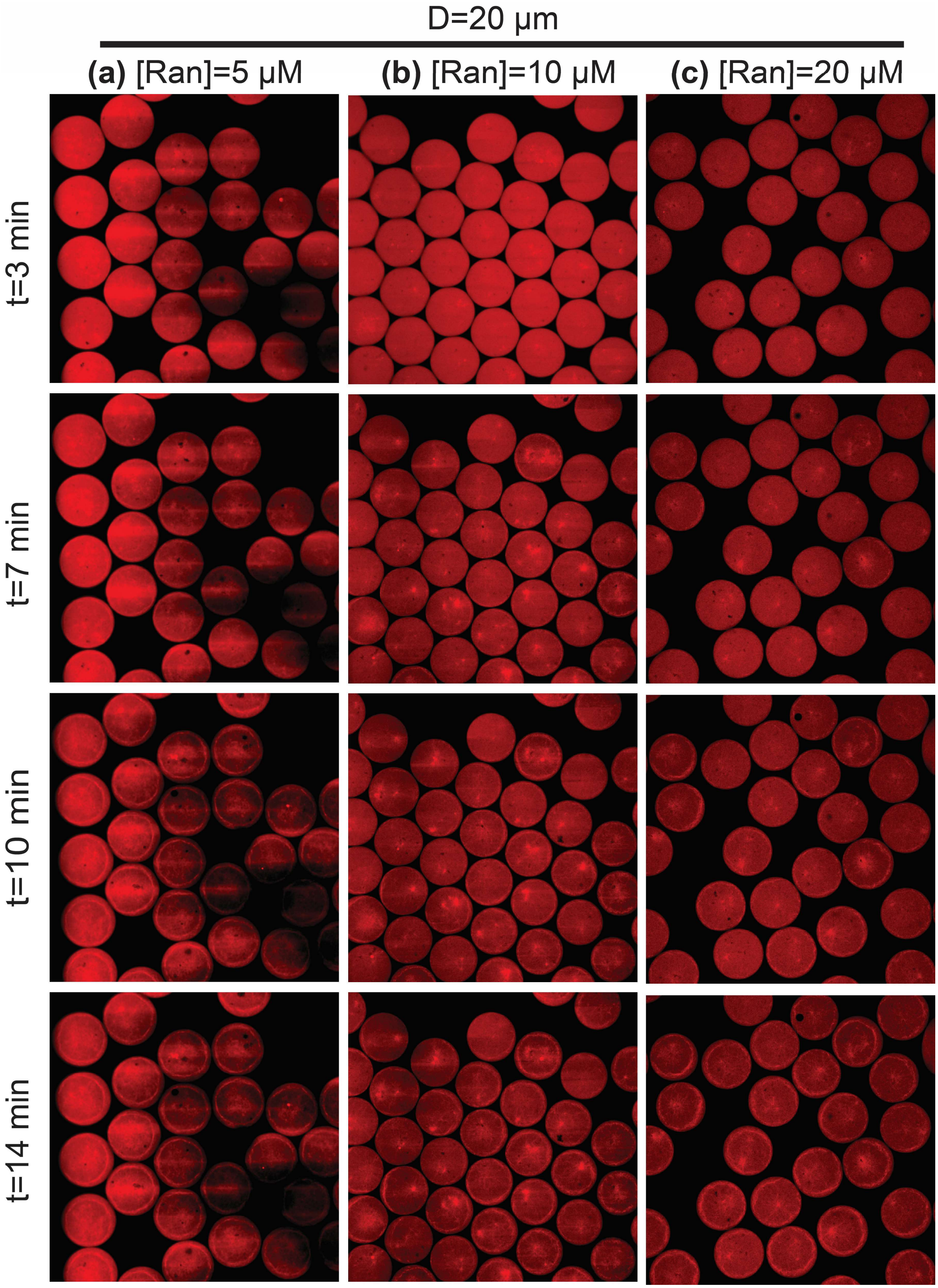

**Figure S4. Wide-field images showing the assembly of flower structures**

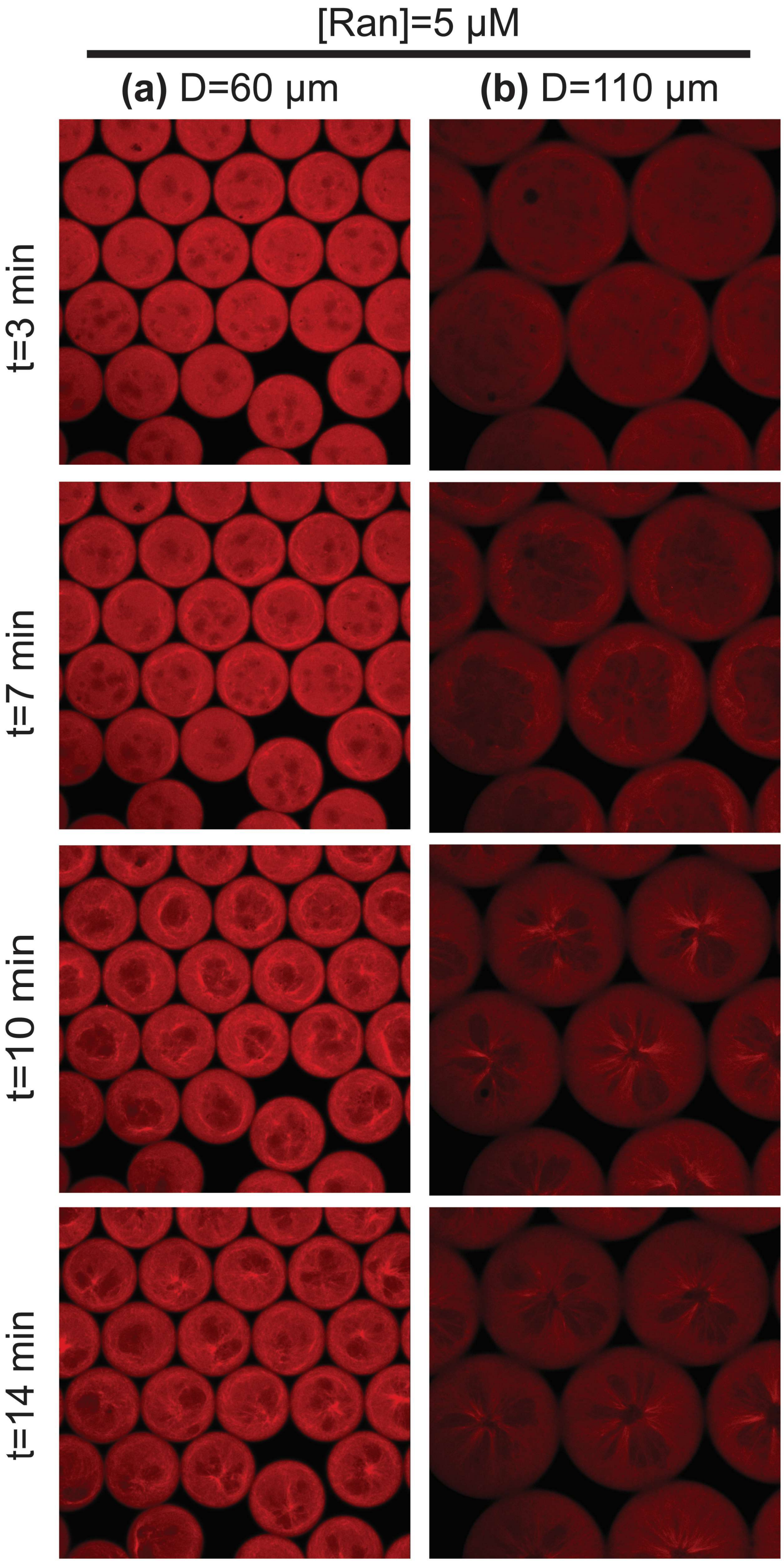

**Figure S5. Number of partitions in flower-shaped MT networks**

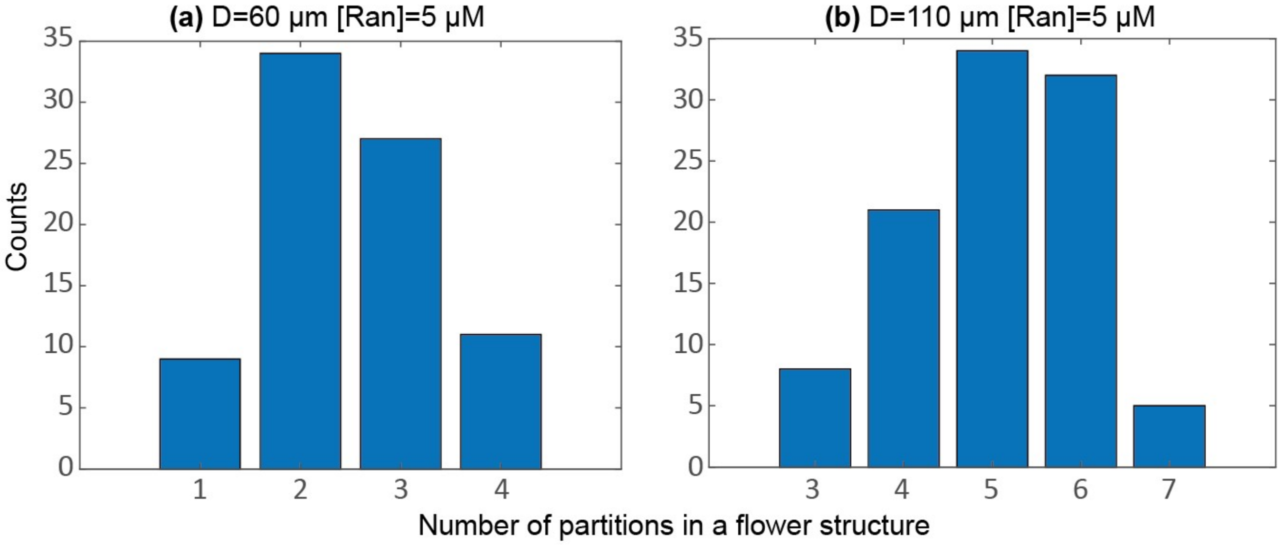


Number of partitions or void areas in flower-shaped MT networks. (a) The droplet size is *D=60 μm*. The histogram was generated by tracking a total of *n=80* droplets. The flower structure corresponds to Fig. 3(a) in the main text. (b) The droplet size is *D=110 μm*. The histogram was generated by tracking a total of *n=80* droplets. The flower structure corresponds to Fig. 3(b) in the main text.

**Figure S6. Wide-field images showing the assembly aster structures**

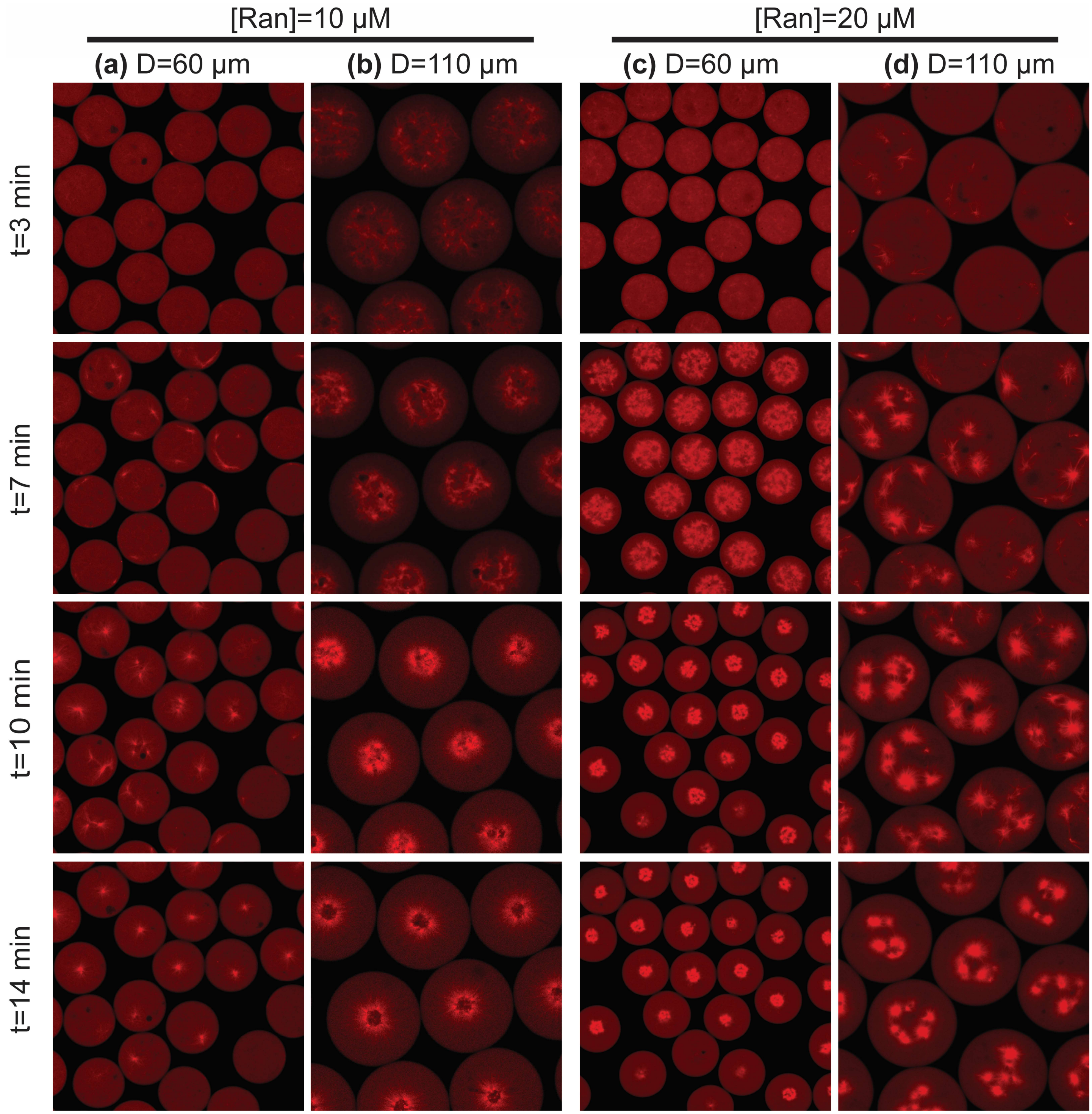

**Supplementary Information Movies**

**Movie S1.** Videos showing ring-shaped MT networks inside droplets with *D=20 μm* and Ran concentrations of 5 μM, 10 μM, and 20 μM.

**Movie S2.** Videos showing flower-shaped MT networks. The test conditions are *D=60 μm* and *Ran=5 μM*, and *D=110 μm* and *Ran=5 μM.*

**Movie S3**. Videos showing the assembly of MT asters. From the left to the right, the test conditions are *D=60 μm* and *Ran=10 μM*, *D=110 μm* and *Ran=10 μM*, *D=60 μm* and *Ran=20 μM,* and *D=110 μm* and *Ran=20 μM.*

**Movie S4.** Widefield videos showing flower-shaped MT networks. Left: *[Ran]=5 μM* and *D= D=60 μm*. Right: *[Ran]=5 μM* and *D=110 μm.*

**Movie S5.** Widefield videos showing MT networks with aster architectures. From the left to the right, the test conditions are *D=60 μm* and *Ran=10 μM*, *D=110 μm* and *Ran=10 μM*, *D=60 μm* and *Ran=20 μM,* and *D=110 μm* and *Ran=20 μM.*

**Movie S6.** Effect of inhibiting motor proteins in confined extracts by adding Vanadate. The final concentration of Vanadate in the droplets is 500 nM. The droplet size is *D=110 μm*, and the Ran concentration is *[Ran]=10 μM.*
